# Creating a bottleneck: Robust LC3B lipidation analysis by adjusting autophagic flux with low concentrations of Bafilomycin A1

**DOI:** 10.1101/2021.07.12.452036

**Authors:** Martina P. Liebl, Sarah C. Meister, Lisa Frey, Kristina Hendrich, Anja Klemmer, Christian Pohl, Viktor Lakics

## Abstract

Autophagic flux can be quantified based on the accumulation of lipidated LC3B in the presence of late-stage autophagy inhibitors. This method has been widely applied to identify novel compounds that activate autophagy. Here we scrutinize this approach and show that bafilomycin A1 (BafA) but not chloroquine is suitable for flux quantification due to the stimulating effect of chloroquine on non-canonical LC3B-lipidation. Significant autophagic flux increase by rapamycin could only be observed when combining it with BafA concentrations not affecting basal flux, a condition which created a bottleneck, rather than fully blocking autophagosome-lysosome fusion, concomitant with autophagy stimulation. When rapamycin was combined with saturating concentrations of BafA, no significant further increase of LC3B lipidation could be detected over the levels induced by the late-stage inhibitor. The large assay window obtained by this approach enables an effective discrimination of autophagy activators based on their cellular potency. To demonstrate the validity of this approach, we show that a novel inhibitor of the acetyltransferase EP300 activates autophagy in a mTORC1-dependent manner. We propose that the creation of a sensitized background rather than a full block of autophagosome progression is required to quantitatively capture changes in autophagic flux.

## Introduction

Modulation of macro-autophagy (hereafter referred to as autophagy) presents a promising therapeutic strategy in a variety of human diseases, ranging from cancer to neurodegeneration. A key pharmacological target in this respect is the mechanistic Target Of Rapamycin (mTOR) complex I (mTORC1) [1, 2, 3]. MTORC1 is a serine/threonine protein kinase complex, which is recognized as a master regulator of cell growth, proliferation, metabolism, and ageing. It negatively regulates autophagy via phosphorylation of the ULK1 complex [4, 5, 6], the PI3K complex [7], as well as the Transcription Factor EB (TFEB) [8]. MTORC1 inhibition by the macrolide rapamycin and its analogs increase autophagy by blocking the inhibitory phosphorylation of these downstream targets. Although several studies confirmed that rapamycin treatment facilitates degradation of protein aggregates in various disease models through autophagy [9, 10, 11, 12, 13, 14] and prolongs life-span in yeast and animal models [15, 16, 17, 18, 19, 20], its wider use as a drug is relatively limited due to strong anti-proliferative and immunosuppressive effects [21, 22]. Instead, rapamycin emerged as a broadly used key tool compound for studying the effects of autophagy stimulation, together with other tool compounds which are inhibitors of the autophagosome–lysosome fusion. The latter process requires an acidic luminal microenvironment in lysosomes maintained by the proton-pumping lysosomal vacuolar H^+^-ATPase (v-ATPase), a multi-subunit protein complex. Two other macrolides, bafilomycin A1 (BafA) and concanamycin A (ConA), inhibit the v-ATPase by binding the membrane-spanning core forming domain (V_0_ subcomplex) of the enzyme [23, 24, 25, 26, 27]. Due to the inhibition of the v-ATPase, BafA collapses the pH-gradient between lysosomes and the cytosol, which subsequently also blocks autophagosome-lysosome fusion. Another modulator of autophagosomal-lysosomal membrane fusion is the anti-malaria drug chloroquine (CQ) which is believed to increase the lysosomal pH by its di-protonation, upon the uptake into the lysosomal lumen [28, 29, 30]. More recently, a direct effect of CQ on lysosomal pH was questioned and suggested that it rather inhibits autophagosome-lysosome fusion without altering lysosome acidity [31]. The late-stage autophagy inhibitors introduced above provide the basis to study an increase of microtubule-associated protein 1 light chain 3B (LC3)-lipidation as a marker for autophagy activation. Although autophagy activation facilitates localization and ATG4-dependent lipidation of cytosolic LC3 to the elongating inner and outer autophagosomal membrane (now termed: LC3-II), using LC3-II levels as a marker of autophagic flux stimulation is only conclusive in the presence of late-stage autophagy inhibition. The *bona fide* autophagy receptor p62/SQSTM1 is another widely used marker in mechanistic studies. P62/SQSTM1 interacts directly with ubiquitinated cargo through its C-terminal UBA-domain, although cargo ubiquitination is not a general prerequisite for p62/SQSTM1 to recognize and bind cargo [32, 33, 34]. In selective autophagy, P62/SQSTM1 bridges autophagic cargo to the autophagosomal lumen by interacting with LC3-II through its LC3-interacting region (LIR) motif [32, 33]. However, LC3-II-independent interaction of p62/SQSTM1 with autophagosomes has also been described [35]. Basal autophagy keeps levels of p62/SQSTM1 relatively low, whereas inhibition of autophagy increases p62/SQSTM1 levels [36, 37]. Monitoring levels of LC3-II in the presence, and levels of p62/SQSTM1 in the absence of late-stage autophagy inhibitors is a widely used practice in autophagy research, and different studies use various late-stage inhibitors at different concentrations, often at saturation to fully inhibit autophagosome-lysosome fusion. Moreover, these compounds are used for various treatment times to capture autophagic flux, making it difficult to find the best starting conditions for a robust autophagic flux assay.

Our goal was to set up a simple but robust method to test novel potential autophagy activators in cell-based experiments, using increased LC3-lipidation as a read-out. An important criterion for such a method is a large assay window, allowing to discriminate between autophagy inducers with different potencies and to reliably discriminate true autophagy activators from agents that impair or block autophagic degradation itself. We have therefore carefully titrated late-stage autophagy inhibitors to find the most suitable concentration applicable in cellular autophagy assays and determined their half-maximal effective concentrations (EC_50_) for the modulation of LC3-II- and p62/SQSMT1 levels, using Western blotting. We demonstrate that, in contrast to applying saturating concentrations of late-stage inhibitors in flux assays, increased LC3-lipidation in response to rapamycin can be best captured in presence of a non-saturating concentration of BafA, which does not inhibit autophagosomal-lysosomal fusion at a basal level. Moreover, high concentrations of BafA or CQ decreased mTORC1 activity, demonstrating additional adverse effects of saturating concentrations. We have also found that CQ was not suitable in autophagic flux assays since it increased LC3-lipidation independently of canonical macro-autophagy. This approach is not only suitable to capture autophagy activation by rapamycin, but also robustly detects increased flux caused by starvation of serum, HBSS starvation, as well as the inhibition of acetyltransferase EP300. We also confirm that EP300 has multiple targets besides mTORC1 since we detected a p62/SQSTM1 decrease through a mechanism downstream or independent of mTORC1. In summary, we present a robust Western-blot based autophagic flux assay with a high assay window suitable to discriminate autophagy activators from late-stage autophagy inhibitors.

## Results

### Non-saturating, but not saturating concentrations of BafA result in a wide assay window when measuring LC3-II lipidation in response to autophagy activation

Clonal cell lines often show cell-type specific sensitivity towards late-stage autophagy inhibitors and longer than 24 h treatment with these compounds frequently results in acute toxicity and cell death, especially at higher concentrations (data not shown). To avoid unnecessary high concentrations of late-stage autophagy inhibitors to measure flux, we sought to determine the lowest saturating concentrations of BafA and CQ in HeLa and SK-N-MC cells, widely used as autophagy models in vitro. We defined a saturating concentration of a late-stage autophagy inhibitor as the lowest concentration which maximally increases LC3-II and/or p62/SQSTM1 levels under basal conditions and therefore represent a full block of the autophagosome-lysosome fusion. To that end, both cell lines were treated with various concentrations of BafA (**Fig. 1A and Suppl. Fig. 1A**) and CQ (**Fig. 1B, Suppl. Fig 1B**). 10 nM BafA treatment of HeLa cells for 24 h resulted in maximal LC3-II increase as detected by Western blot of EC_50_: 5.6 nM. As expected, the increase of LC3-II correlated well with the accumulation of the autophagy receptor p62/SQSTM1, since BafA blocks the degradation of p62/SQSTM1 by basal autophagy. The EC_50_ of 6.5 nM for p62/SQSTM1 accumulation was nearly identical to the half-maximal concentration required for LC3-II increase. The EC_50_ values for LC3-II and p62/SQSTM1 in SK-N-MC were very similar at 7.5 nM and 8.5 nM, respectively **(Suppl. Fig. 1A)**. Based on these measurements, we regarded 10 nM BafA as the lowest saturating concentration, used as such in subsequent experiments. In contrast, 2.5 nM BafA only minimally increased LC3 lipidation or p62/SQSTM1 accumulation in both cell lines, indicating that at basal conditions autophagic flux is not strongly impaired in presence of this low dose of BafA. Therefore, with respect to inhibition of autophagosomal-lysosomal fusion, we called 2.5 nM a “non-saturating concentration” of BafA.

**Figure 1.**
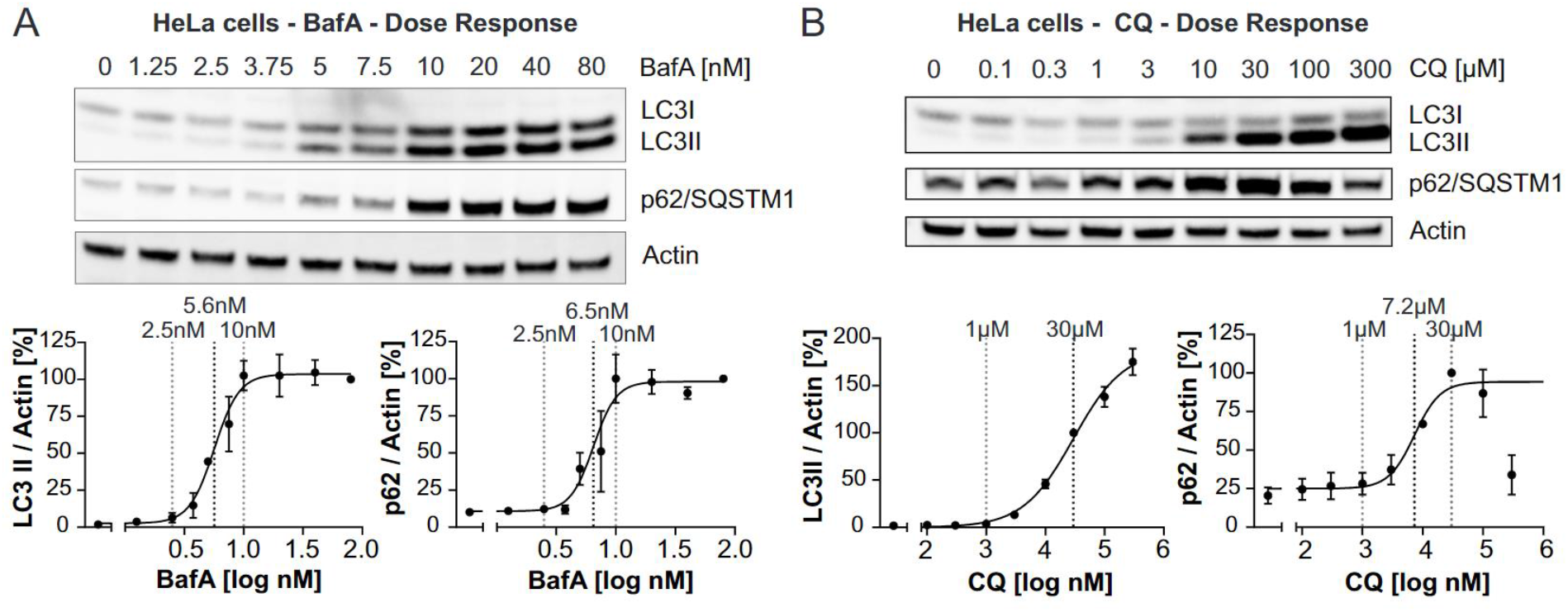
Determination of saturating concentrations of BafA and CQ by Western blotting in HeLa cells. (**A**), (**B**) Treatment of HeLa cells for 24 h with various concentrations between 1.25 and 80 nM BafA or 0.1 – 300 μM CQ. Top: representative Western blots for LC3, p62/SQSTM1 and actin; bottom: quantifications from 3 independent experiments (mean +/− SEM). For quantification, LC3-II levels at each concentration were normalized to the LC3-II levels at the highest concentration applied. (**A**) BafA increased LC3-II levels with an EC_50_ of 5.6 nM and p62/SQSTM1 levels with an EC_50_ of 6.5 nM. (**B**) For CQ, the calculated EC_50_ for LC3-II and p62/SQSTM1 increase were 30 μM and 7.2 μM, respectively. For curve fitting, a non-linear regression curve fit (log(inhibitor) vs. response-variable slope (four parameters)) was applied.

In contrast to BafA, when CQ was used, a saturating concentration for the LC3-II increase could not be established at non-toxic concentrations in any of the two cell lines. The highest concentration of CQ applied to HeLa cells (300 μM) strongly decreased cell confluency and changed cellular morphology, indicating toxicity (data not shown). However, even at 300 μM, the CQ induced LC3-II increase was not plateaued in HeLa cells **(Fig. 1B)**. Similarly, in SK-N-MC cells no saturation of LC3-II has been observed either (established up to 100 μM CQ, **Suppl. Fig. 1B)**, in contrast to p62/SQSTM1 accumulation, which reached a plateau at 30 μM CQ applied for 24 h (HeLa: EC_50_: 7.2 μM; SK-N-MC: EC_50_: 9 μM), indicating that basal autophagic flux was fully blocked at this concentration (**Fig. 1B and Suppl. Fig. 1B**). Therefore, we considered 30 μM CQ a “saturating concentration” regarding p62/SQSTM1 accumulation. Similar to 2.5 nM BafA, no significant increase of LC3-II or p62/SQSTM1 was observed in the presence of 1 μM CQ, therefore we used 1 μM CQ as a non-saturating concentration in further experiments. To avoid potential toxicity caused by long incubation times with late-stage autophagy inhibitors, we also tested the effect of blocking autophagosome-lysosome fusion only during the last 2 h of rapamycin-treatment. In these conditions, BafA-induced LC3-II and p62/SQSTM1 levels were maximal at around 500 nM and again, CQ-induced LC3-II levels did not reach a plateau up to 300 μM (**Suppl. Figs. 2A and B**). P62/SQSTM1 accumulation in response to CQ was saturated at 100 μM.

To find conditions allowing to capture the LC3-II increase by rapamycin with an optimal assay window, we performed a time course experiment, treating HeLa cells for different durations with combinations of either vehicle or rapamycin in the presence of selected saturating or non-saturating concentrations of BafA (**Fig. 2A**). Although we observed earlier that 10 nM BafA was saturating when applied for 24 h, we also included a much higher concentration (2 μM), which was tolerated by the cells only for up to 8 h, to test complete inhibition of autophagosome-lysosome fusion at shorter treatment durations. In additional experiments we have confirmed that 2 μM BafA was saturating in cells treated for 2 h (**Suppl. Fig. 2A**). Based on the accumulation of p62/SQSTM1, the time course experiment confirmed the inhibition of autophagy by both 10 nM and 2 μM BafA, whereas there was no p62/SQSTM1 accumulation observed at any time point when cells were treated with 2.5 nM BafA. A robust p62/SQSTM1 increase was observed at both saturating concentrations as early as 8 h (**Fig. 2B, left panel**). In contrast, rapamycin co-treatment only showed an additional increase of LC3-II over the BafA only condition, when it was combined with the non-saturating concentration of 2.5 nM BafA. A robust LC3-II increase was observed after 16 h, but not at earlier time points, and became even more pronounced after 24 h treatment (**Fig. 2A, right panel**).

**Figure 2.**
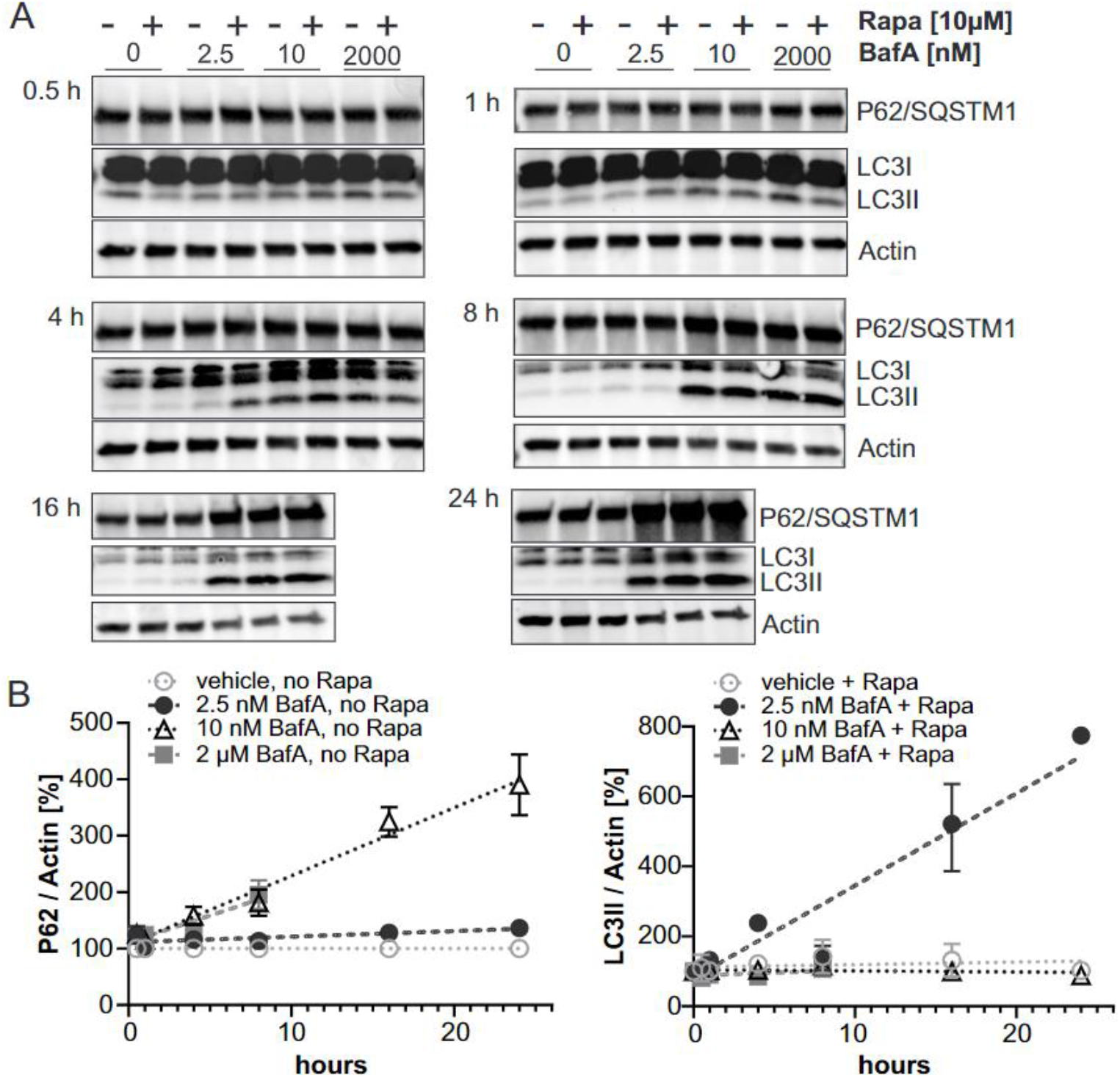
Determination of the optimal assay window for capturing LC3-II increase in response to rapamycin. (**A**) HeLa cells were co-treated for indicated time intervals (0.5 – 24 h) with rapamycin or vehicle in the presence or absence of 2.5 nM, 10 nM or 2 μM (the latter only tolerated for up to 8 h) BafA. Representative Western blots of LC3-II, p62/SQSTM1 and actin are shown for each treatment time point. (**B**) Quantification of Western blots from 2 independent experiments (mean +/− SEM; linear fit). P62/SQSTM1 levels (left panel) from cells treated with BafA in the absence of rapamycin were quantified, normalized to actin, and subsequently compared to p62/SQSTM1 levels treated with DMSO only (0 nM BafA). LC3-II levels (right panel) were normalized to actin. The LC3-II increase caused by 10 μM Rapamycin over time in the presence of either 0, 2.5, 10 or 2000 nM BafA is compared to the levels induced by the respective BafA-concentrations without rapamycin which was set to 100 %.

Based on these time course experiments, we chose 24 h treatment as most suited to capture autophagy activation with rapamycin in the presence of late-stage autophagy inhibition. We performed additional experiments to detect rapamycin-increased LC3 lipidation using co-treatment with the lowest saturating concentration of late-stage autophagy inhibitors. Contrary to previously published work, in both HeLa and SK-N-MC cells, we were not able to show any significant additional LC3-II accumulation above the level caused by the late-stage-inhibitor alone in response to rapamycin treatment when a saturating concentration of BafA was used (**Fig. 3A, Suppl. Fig. 1C, Suppl. Fig. 2C**). In the same experiments, mTOR S2448 phosphorylation was decreased when saturating concentrations of rapamycin were applied (**Suppl. Fig. 2E**). A strong, statistically significant increase in LC3 lipidation could only be detected when rapamycin was combined with a non-saturating concentration of BafA (i.e. 2.5 nM). Furthermore, we were also not able to capture any LC3-II increase by rapamycin if a saturating concentration of BafA was applied only during the last 2 h of rapamycin treatment (**Suppl. Fig. 2C**). Since Western blotting experiments can show relatively high variability, we routinely used rapamycin as a positive control in all experiments aimed to test autophagy stimulation, in the presence of 2.5 nM BafA. Under these conditions, in 86 % of 35 independent experiments performed by 3 different operators, rapamycin increased LC3-II levels by more than 2-fold (**Fig. 3B**). Hence, in our subsequent autophagy flux experiments, results were only considered to be valid if at least 2-fold increase of LC3-II by rapamycin was observed. Generally, the observed assay window of rapamycin-induced LC3-II increase is 2 to 10-fold over the BafA induced levels which makes the assay suitable to compare the cellular potency of different autophagy activators within the same experiment, when analyzed side-by-side in a Western blotting experiment. This suggests that when treating clonal cell lines for 24 h with autophagy modulators – a frequent practice in the literature – a non-saturating concentration of BafA is more suitable than a saturating concentration to capture an autophagy-activating effect of a potential inducer.

**Figure 3.**
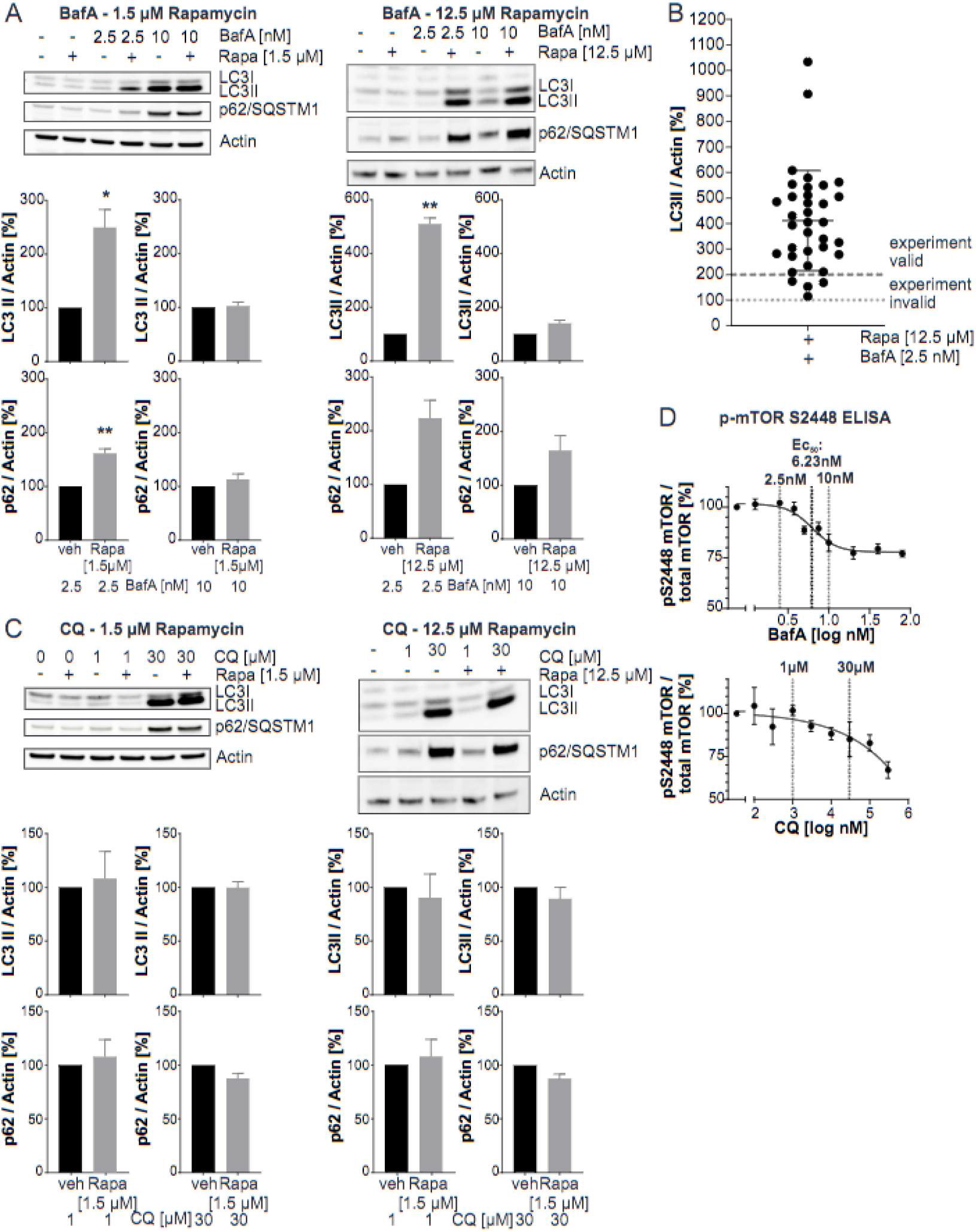
Rapamycin-treatment combined with saturating and non-saturating concentrations of autophagosome-lysosome fusion inhibitors. (**A**) Representative Western blots for LC3-II, p62/SQMST1 and actin (top) and quantifications (n= 3-4; mean +/− SEM) of HeLa cells co-treated with 1.5 μM (left panel) or 12.5 μM (right panel) rapamycin and 2.5 nM or 10 nM BafA for 24 h. Statistical comparison: one sample t-test, * indicates p ≤. 0.05; ** indicates p ≤ 0.01. (**B**) Compiled quantification of Western blots detecting LC3-II increase by 12.5 μM rapamycin in HeLa cells co-treated with 2.5 nM BafA for 24 h normalized to LC3-II levels in HeLa cells treated with 2.5 nM BafA for 24 h only (100 %). The quantification was compiled from 35 independent biological experiments (n = 35) performed by 3 different operators. (**C**) Representative Western blots for LC3-II, p62/SQMST1 and actin (top) and quantifications (n= 3; mean +/− SEM) of HeLa cells co-treated with 1.5 μM (left panel) or 12.5 μM (right panel) rapamycin and 1 μM or 30 μM QC for 24 h. Statistical analysis: one sample t-test; no significant differences were observed. (**D**) Lysates from HeLa cells treated with a dose range of BafA (1.25 nM – 80 nM) or CQ (0.1 μM – 30 μM) for 24 h were analyzed with a mTOR MSD ELISA assay multiplexing total mTOR and pS2448 mTOR. For BafA, an EC_50_ of 6.23 nM with a saturating effect at 20 % signal decrease was determined. Curves represent one biological replicate with 3 technical replicates (n =1; technical mean +/− SEM). For curve fitting, a nonlinear regression curve fit (Sigmoidal, four parameter logistic, X is log(concentration)) was applied.

To compare the suitability of BafA with those of CQ to capture LC3-II increases by autophagy activators, HeLa and SK-N-MC cells were treated with rapamycin in the presence of a saturating and a non-saturating concentration of CQ (**Fig. 3C**, **Suppl. Fig 2D**). In HeLa cells neither a saturating, nor a non-saturating concentration of CQ was sufficient to capture any rapamycin-induced LC3-II increase (**Fig. 3C, Suppl. Fig. 2D**). In contrast to HeLa cells, we found that in SK-N-MC cells it was possible to capture a significant increase of LC3-II by rapamycin over a non-saturating concentration of CQ. (**Suppl. Fig. 1C**). However, similar to HeLa cells, saturating CQ concentrations also failed to help the detection of LC3-II increases by rapamycin with a usable effect size. Taken together, these data underline that saturating concentrations of late-stage autophagy inhibitors are not suited to capture autophagy enhancing effects of rapamycin in these cell lines.

Our subsequent experiments were aimed towards explaining the lack of LC3-II increase in response to rapamycin in presence of a saturating concentration of BafA. Since mTORC1 activity depends on its localization to lysosomal membranes, we tested how mTOR activity is modulated by BafA-or CQ-dependent effects on lysosomes. To investigate whether a saturating concentration of BafA or CQ induced a negative feedback loop and impaired early autophagy activation in an mTORC1-dependent manner, we checked using an MSD-ELISA whether BafA and CQ increase mTOR S2448 phosphorylation (**Fig. 3D**). Rapamycin served as a positive control and reduced mTOR S2448 phosphorylation by more than 50 % in the same experiment (*not shown*), corresponding to mTOR inhibition and autophagy activation. However, neither BafA nor CQ increased mTOR S2448 phosphorylation, so this could not explain why in presence of saturating concentrations of late-stage autophagy inhibitors no additional LC3-II increase with rapamycin was detected. Instead of an increase, a concentration-dependent decrease of mTOR S2448 phosphorylation was observed with late-stage autophagy inhibitors. BafA decreased mTOR S2448 phosphorylation with an EC_50_ of 6.23 nM and reached a maximum reduction of 25% (**Fig. 3D**). Interestingly, this EC_50_ was in line with the EC_50_ values determined for BafA-induced LC3-II and p62/SQSMT1 accumulation, suggesting a direct link between impairment of autophagosome-lysosome fusion and mTOR inhibition. For CQ, no saturating concentration and no EC_50_ with respect to mTOR S2448 phosphorylation could be determined at non-toxic concentrations. For both late-stage autophagy inhibitors no decrease of mTOR S2448 phosphorylation was observed in the presence of non-saturating concentrations (2.5 nM BafA; 1 μM CQ). However, if BafA or CQ at saturating concentrations impaired the sensitivity of the autophagic system towards autophagy activators, this inhibitory effect must have occurred either downstream or independently of mTOR activation.

### Non-saturating BafA concentrations discriminate autophagy activators from late-stage inhibitors

Based on our observation that rapamycin only increased the LC3-II levels in the presence of a non-saturating concentration of BafA, we were concerned that under non-saturating conditions an autophagosome-lysosome fusion inhibitor would behave similarly to an autophagy activator due to the potential additive effect of both compounds on the LC3-II increase. To further investigate whether non-saturating or saturating concentrations of BafA could discriminate early-stage autophagy activators from late-stage autophagy inhibitors, multiple concentrations of CQ and ConA were combined with a non-saturating BafA concentration (**Figs. 4A and B**). Indeed, while 2.5 nM BafA alone did not block basal fusion of autophagosomes with lysosomes and no LC3-II accumulation could be observed, CQ increased LC3-II levels equally in the presence and in the absence of BafA (**Fig. 4A**), clearly identifying it as a late-stage inhibitor. Interestingly, the LC3-II increase by CQ was even stronger in the absence than in the presence of BafA, confirming that CQ-induced single membrane lipidation requires v-ATPase activity [39]. P62/SQSTM1 levels were also increased by CQ in the absence of BafA, further justifying the classification of similarly behaving test compounds in the autophagic flux assay as potential late-stage autophagy inhibitors. ConA (**Fig. 4B**) similarly increased LC3-II levels regardless of the presence or absence of a non-saturating BafA concentration. Moreover, in the presence of 2.5 nM BafA, 250 pM ConA was sufficient to increase LC3-II levels to the extent observed only at 500 pM ConA when BafA was absent. This additive effect on the LC3-II increase is expected when applying two late-stage autophagy inhibitors at non-saturating concentrations. The LC3-II increase in the presence and absence of BafA strongly correlated with increased levels of p62/SQSTM1, again supporting the classification of similarly behaving compounds as late-stage autophagy inhibitors. Taken together, applying a non-saturating of BafA in the flux assay allows to discriminate autophagy activators from compounds blocking autophagosomal-lysosomal fusion if the appropriate control, namely the application of the compound at multiple concentrations also in the absence of BafA, is included. Considering changes in p62/SQSTM1 levels induced by a test compound in the absence of BafA will further help to discriminate autophagy activators from late-stage autophagy inhibitors.

**Figure 4.**
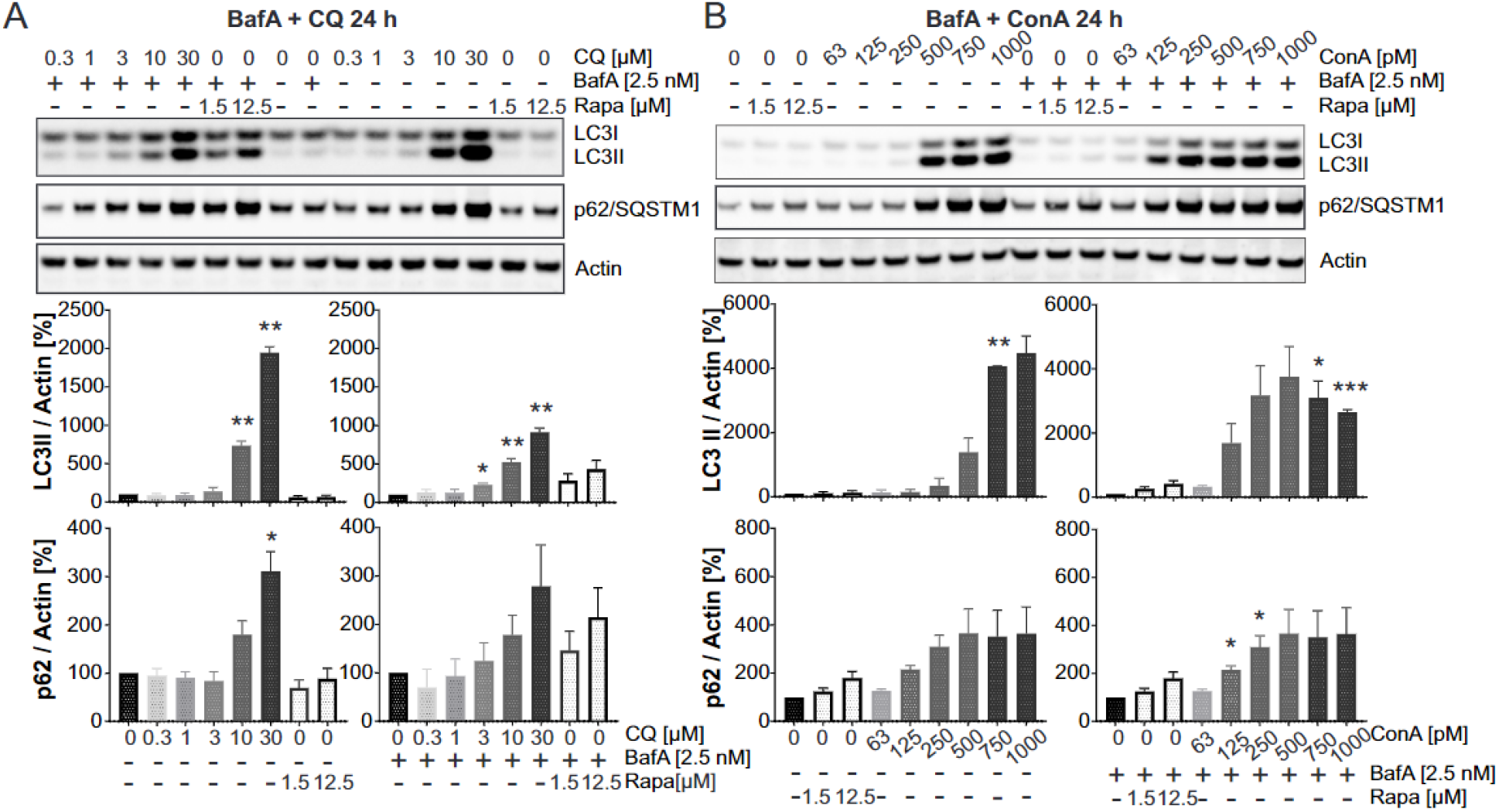
Co-treatment of HeLa cells with BafA and additional autophagosome-lysosome fusion inhibitors. (**A**) Representative Western blots for LC3, p62/SQSTM1 and actin (top) and quantification (n=3; mean +/− SEM) of HeLa cells co-treated with either DMSO (left) or 2.5 nM BafA (right) and a dose range (0.3 – 30 μM) of CQ for 24 h. Statistics: One sample t-test, compared to 100 % control (either DMSO only or 2.5 nM BafA only). (**B**) Representative Western blots for LC3, p62/SQSTM1 and actin (top) and quantification (n=3; mean +/− SEM) of HeLa cells co-treated with either DMSO (left) or 2.5 nM BafA (right) and a dose range (63 pM – 1 nM) of ConA for 24 h. Statistics: One sample t-test, compared to 100 % control (either DMSO only or 2.5 nM BafA only).

### Autophagic flux analysis after serum and HBSS starvation in the presence of non-saturating concentrations of BafA

A robust flux assay with a large window is expected to capture LC3-II increase by various stimulators, including serum withdrawal (applying HBSS as cell culture medium) or serum starvation in the presence of a non-saturating concentration of BafA. To test whether reduction of serum (fetal bovine, FBS) levels in the medium from 10 % to 1.5 % activated autophagy, LC3-II levels were measured 4 h and 24 h after reduction of the FBS concentration (**Fig. 5A**). 4 h as well as 24 h of serum starvation strongly increased LC3-II levels in the presence, but not in the absence, of a non-saturating concentration of BafA. Combining serum reduction with maximal rapamycin concentration had an additive effect on the LC3II-level increase, suggesting that serum depletion activated autophagy has an mTORC1-independent component. Moreover, combining 3 hours of HBSS starvation (no serum, no amino acids, glucose is present in HBSS) with 2.5 nM BafA treatment allowed to capture a strong increase in LC3-II levels in comparison to non-starved cells treated with 2.5 BafA (**Fig. 5B**).

**Figure 5.**
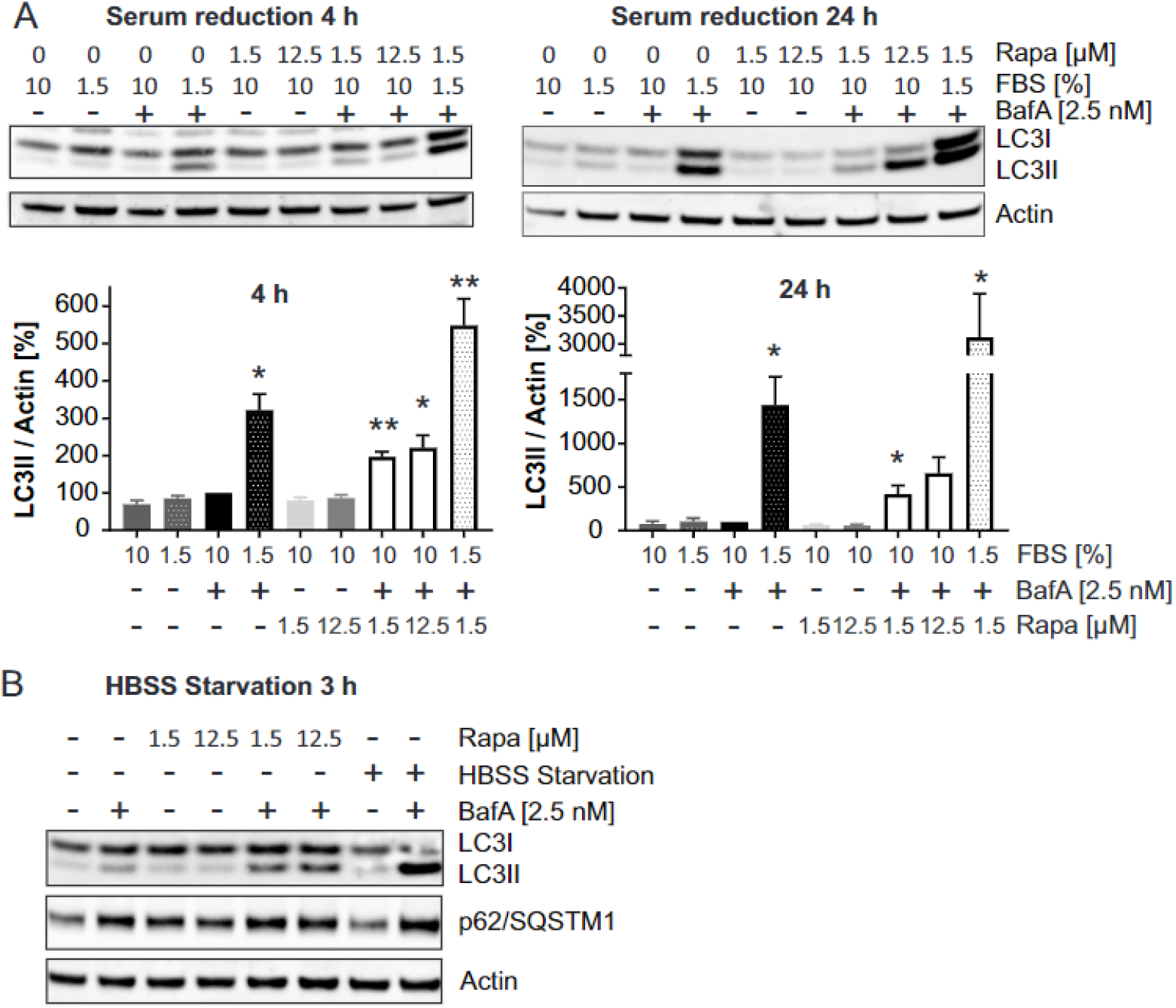
Capturing autophagic flux increase through serum starvation and HBSS starvation. (**A**) HeLa cells were kept in standard medium (10 % FBS) or in FBS-reduced medium (1.5 %) for 4 or 24 h. Medium change was combined with (co)-treatment of cells with 2.5 nM BafA and / or 1.5 or 12.5 μM rapamycin. Shown are representative Western blots of LC3 and actin (top) as well as quantifications (n = 3; mean +/− SEM; bottom). Statistics: One-sample t-test, compared to control (2.5 nM BafA). (**B**) HeLa cells were kept in standard cultivation medium or HBSS for 3 h in the presence or absence of 2.5 nM BafA and / or 1.5 μM or 12.5 μM rapamycin. Shown is a representative Western blot for LC3, p62/SQSTM1 and actin.

### Evaluation of EP300 inhibitors using non-saturating concentrations of BafA

To further extend the applicability of our approach to the detection of autophagy modulation, we investigated whether the inhibition of the acetyltransferase EP300 impacts autophagic flux, as expected. To that end, we tested a novel, potent EP300 inhibitor synthesized in AbbVie (compound 1) [38], which binds to the histone-acetyltransferase-domain of EP300, in our autophagy flux assay (**Fig. 6A**). Similar to rapamycin, compound 1 increased LC3-II levels in the presence, but not in the absence of BafA (**Fig. 6B**). Moreover, in the absence of BafA, EP300 inhibition strongly decreased levels of p62/SQSTM1 (**Fig. 6C**), another indication of a potential stimulatory effect on autophagy. To reveal the mechanism by which the compound stimulates autophagy, we analyzed the phosphorylation status of the two mTORC1 effectors p70^S6^ kinase (p70^S6k^) and S6 ribosomal protein which was significantly decreased by compound 1 (**Fig. 6C**).

**Figure 6.**
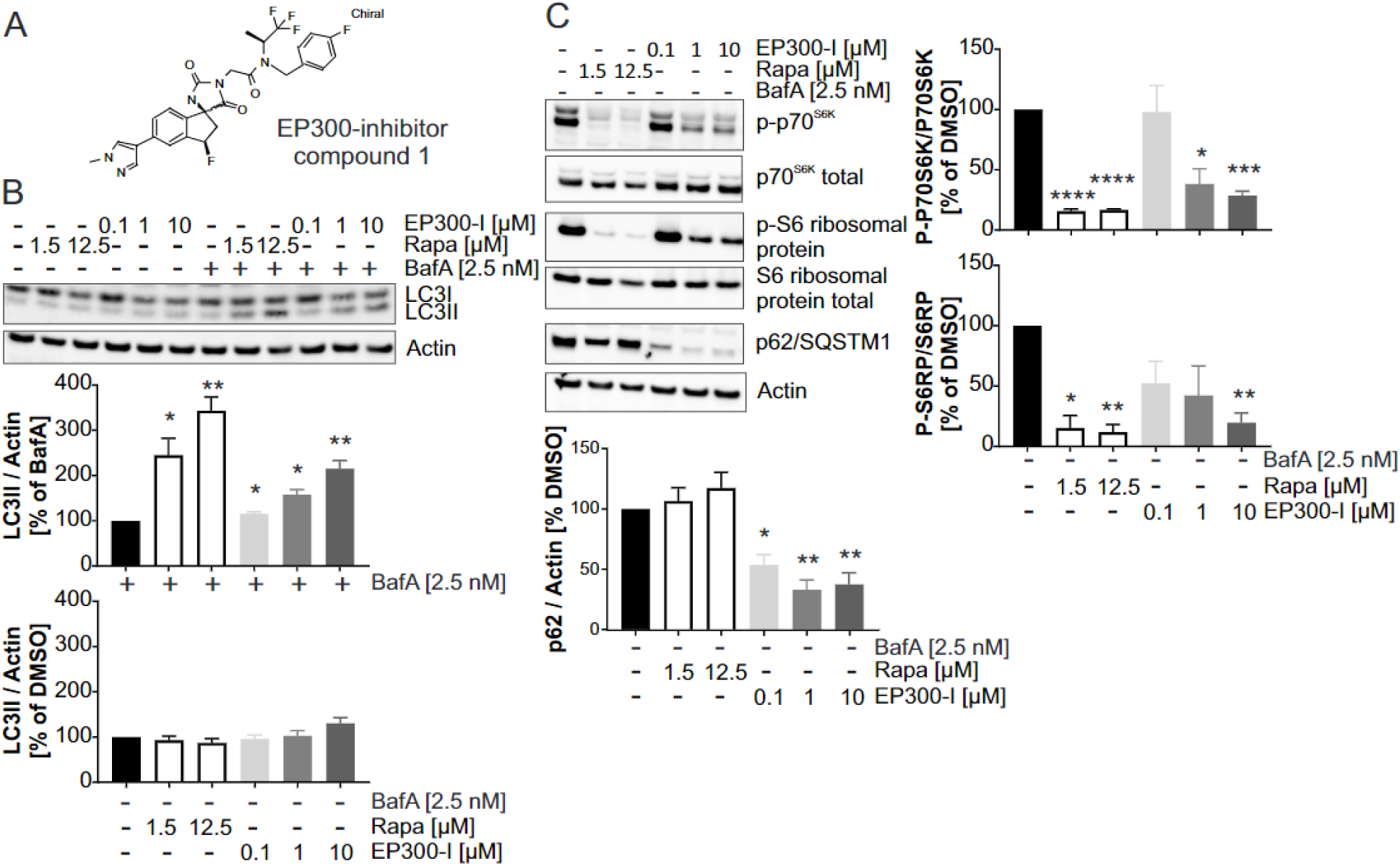
Demonstrating the autophagy stimulating effect of a novel acetyltransferase EP300 inhibitor. (**A**) Chemical structure of the HAT-domain binding EP300 inhibitor (compound 1) which was applied to HeLa cells for 24 h. (**B**) HeLa cells were treated co-treated with a dose range of an EP300-inhibitor (0.1 – 10 μM) and 2.5 nM BafA for 24 h. Co-treatment of cells with rapamycin (1.5 μM and 12.5 μM) and 2.5 nM BafA was applied as a positive control. Shown is a representative Western blot for LC3 and actin (top) as well as a quantification (bottom) from Western blots of 4 independent biological replicates (n = 4; mean +/− SEM). Statistics: One-sample t-test; comparison to 100 % control (either 2.5 nM BafA – upper panel; or DMSO – lower panel). (**C**) HeLa cells were treated with 0.1, 1 or 10 μM of an EP300-inhibitor. Rapamycin (1.5 μM, 12.5 μM) was applied as a control. Shown is a representative Western blot for p-p70SK6, total p70S6K, p-S6 ribosomal protein, total S6 ribosomal protein, p62/SQSTM1 and actin (top), as well as quantifications from Western blots of 4 independent biological replicates (n = 4; mean +/− SEM). Statistics: One-sample t-test compared to 100 % control (DMSO).

## Discussion

One of the most frequently used methods to detect changes in autophagic flux is the Western blot-based, semiquantitative measurement of LC3-II levels upon stimulation or inhibition of macro-autophagy [40]. Most of the current approaches include the use of a late-stage inhibitor of autophagy in a saturating concentration to block lysosomal-autophagosomal fusion, thereby potentially increasing the amount of LC3-II captured by the assay. Although both inhibitors are used interchangeably in autophagic flux assays, we found that BafA, but not CQ, was suitable to capture the increase of LC3 lipidation by the mTORC1 inhibitor rapamycin in two different cell lines. As opposed to the use of BafA, our results showed the lack of saturation for LC3-II levels when cells were treated with various concentrations of CQ, in contrast to the levels of the basal autophagy substrate P62/SQSTM1, which reached saturation. This strongly suggests that CQ, besides blocking autophagosome-lysosome fusion, stimulates unconventional LC3 lipidation on single membrane intracellular compartments, a phenomenon reported earlier [41, 39, 42]. Single membrane LC3 lipidation has been observed in the context of phagocytosis (LC3-associated phagocytosis, LAP), endocytosis (LC3-associated endocytosis, LANDO), micropinocytosis and entosis [41, 43, 44]. Interestingly, we found that CQ increased LC3-II to an even lesser extent in presence the of BafA, although an additive effect was expected when combining two late-stage autophagy inhibitors. A possible explanation could be that the presence of BafA compromises v-ATPase activity, which is indispensable for unconventional LC3-lipidation, resulting in weaker CQ-dependent LC3-II increase when combining CQ and BafA [39, 45]. There are significant differences regarding the molecular mechanisms of these two late-stage inhibitors. Although BafA decreases the lysosomal pH by v-ATPase inhibition, to date it is not fully elucidated at the molecular level how autophagosome-lysosome fusion is inhibited in response to the loss of the pH-gradient between lysosomes and the cytosol [46, 47, 48]. Based on our findings and published data, caution needs to be taken when comparing studies applying different late-stage autophagy inhibitors in autophagy flux assays. We conclude that BafA and CQ cannot be used interchangeably, and at least in the often-used HeLa and SK-N-MC clonal cell line models, CQ is not a suitable late-stage autophagy inhibitor to be applied in an assay using the LC3-II lipidation readout.

BafA was reported to reversibly bind to the V_0_ subunit of v-ATPase and in our hands inhibited the basal autophagic flux with an EC_50_ around 6 nM. While no remarkable increase in LC3-II or p62/SQSTM1 by 2.5 nM BafA treatment was observed at basal conditions, the inhibition curve plateaued already at 10 nM BafA. This represents a very narrow concentration-response similarly to data reported by others [26, 49, 50]. The EC_50_ values or effective concentration ranges in these studies are in line with our measurements, and together with the narrow concentration response, emphasize the need to determine the most suitable concentrations and treatment durations for late-stage inhibitors in each cellular model systems, when performing flux experiments.

Based on previous publications, we expected that completely blocking the autophagosomal-lysosomal fusion with a saturating concentration of BafA would allow to capture maximum LC3 lipidation by rapamycin-induced mTORC1 inhibition. Applying the lowest saturating concentration of BafA to HeLa or SK-N-MC cells strongly increased LC3-II levels, suggesting that in both cell lines there was a high basal autophagic turnover. In contrast, we could not further enhance the strong LC3-II increase observed in the presence of a saturating concentration of BafA by adding rapamycin at any time point investigated in HeLa or SK-N-MC cells. However, a non-saturating concentration of BafA captured a two- to tenfold increase of LC3 lipidation by rapamycin highly reproducibly when applied for at least 16 hours. This assay window is appropriate for comparing the efficacy of potential autophagy modulators in a cellular autophagy flux assays setting, using a Western blot-based LC3-II readout without the need to perform time course experiments. We speculate that the complete blockage of autophagosome-lysosome fusion triggers a negative feedback mechanism, desensitizing the autophagic machinery towards stimulation with autophagy activators. Physiologically, such a mechanism could be relevant to protect cells from excessive autophagosome generation in conditions where autophagosome-lysosome fusion is (transiently) impaired. A significant cross-talk between mTORC1 and lysosomes takes place and mTORC1 directly regulates lysosomal biogenesis by phosphorylating TFEB. TFEB, a master regulator of lysosomal biogenesis, regulates the transcription of multiple lysosomal genes including v-ATPase, directly linking mTORC1 to the transcriptional regulation of autophagy [50; 8]. Moreover, mTORC1 activation requires its localization to the lysosome, ensuring inhibition of autophagy under nutrient rich conditions [51, 52, 53, 54]. Accordingly, we found that long-term treatment with BafA or CQ decreased mTOR S2448 phosphorylation, indicating that late-stage autophagy inhibition promotes mTOR inhibition. In line with this, recently it was also shown that BafA, as well as CQ reduce the lysosomal localization of mTORC1[55] and that v-ATPase inhibition negatively impacts the strong activation of mTORC1 in response to high amino acid concentrations [56]. Noteworthy, the EC_50_ values for the BafA-dependent mTOR inhibition, LC3-II accumulation and p62/SQSTM1 increase were nearly identical (6.2 nM, 5.6 and 6.5 nM, respectively). This reflects the inhibition of the same cellular target, namely v-ATPase and can account for BafA-induced mTOR inhibition if re-localization is directly linked to v-ATPase inhibition. This is in line with a previous report showing that basal mTORC1 activity depends on normal lysosomal function and that late-stage autophagy inhibitors activate early stages of autophagy [57]. Of note, no impact on the mTOR S2448 phosphorylation status was observed with a non-saturating concentration of BafA used for the establishment of the autophagic flux assay described here, further supporting the rationale of avoiding saturating concentrations.

Applying non-saturating concentrations of BafA captured not only autophagy activation by rapamycin, but also the autophagy stimulatory effect of serum withdrawal (HBSS) and serum starvation. Serum starvation activated autophagy is at least partially independent or downstream of mTORC1, as suggested by the additive effect of rapamycin on LC3 lipidation. We did not observe a shutdown of autophagy upon prolonged serum starvation based on LC3-lipidation captured in the presence of BafA, up to 24 h. Previous studies showed a peak of autophagosome-lysosome fusion after 4 h of combined serum / glutamine starvation and reactivation of mTORC1 followed by attenuation of autophagy and lysosomal restoration upon prolonged starvation for 8-12 h [58, 59]. This is not necessarily in contrast with our findings for two reasons: first, previous reports combined serum starvation with the withdrawal of glutamine, which increases intracellular leucine concentrations required for mTORC1 reactivation upon prolonged starvation [60, 59]. Glutamine, as well as a low serum concentration, were still present in our experimental setting, suggesting that the presence of glutamine might be sufficient preventing mTORC1 reactivation in response to prolonged serum starvation. Second, it was shown that mTORC1 reactivation depends on intact lysosomal degradation and that amino acids generated upon prolonged starvation mediate subsequent mTORC1 reactivation [58]. Therefore, although BafA levels were relatively low in our experimental setup, there is a possibility that mTORC1 reactivation in response to prolonged serum starvation was suppressed by lysosomal inhibition.

Nutrient starvation, the most common physiological activator of macroautophagy, depletes the cellular pool of Acetyl-Coenzyme A and therefore facilitates deacetylation of hundreds of proteins [61]. The acetyltransferase EP300 was shown to target the transcription factors NF-k-B, FOXO1 and P53, all part of the transcriptional regulation of autophagy [62, 63, 64], as well as core proteins of the autophagic machinery, namely LC3, Beclin1, ATG5, ATG7 and ATG12 [65, 66, 67]. EP300 directly counteracts deacetylation by sirtuin-1 (SIRT1), a deacetylase implemented in autophagy activation and longevity [68]. In addition, EP300 knockdown, as well as its inhibition with the small molecule c646, have been shown to decrease levels of p62/SQSTM1 as well as mTOR downstream effector phosphorylation, the latter indicating a mTOR-dependent mechanism of autophagy activation [61, 69, 70]. To test whether EP300 inhibition indeed modulates autophagy in the assay setting described herein, we used a novel AbbVie-internal EP300 inhibitor (compound 1) [38]. The increase in LC3 lipidation captured in presence of BafA, together with a strong p62/SQSTM1 decrease observed in the absence of BafA, suggested that EP300 inhibition with compound 1 activated autophagy. Autophagy activation by compound 1 was, as previously described, at least partially mediated via mTOR-inhibition, since EP300 inhibition resulted in significant dephosphorylation of S6 ribosomal protein and p70^S6k^ [61]. Our findings regarding the mTORC1-dependency of the autophagy activating effect of EP300 inhibition are supported by a recent study identifying a lysine residue in the mTORC1 subunit raptor as an acetylation target for EP300 [69]. Moreover, EP300 inhibition, besides increasing LC3 lipidation, strongly decreased p62/SQSTM1 levels. This indicates that mTORC1 inhibition alone is likely not sufficient to drive autophagy-dependent degradation of p62/SQSTM1, whereas EP300 inhibition facilitated p62/SQSTM1 degradation though a mechanism downstream and/or independent of mTORC1. This is of special interest in the light of the recent finding that not LC3-lipidation, but LC3-deacetylation is crucial for its interaction with p62/SQSMT1, and EP300 inhibition facilitated binding of p62/SQSTM1 to LC3 [67]. This suggests that EP300 inhibition likely activated autophagy through both mTORC1 inhibition and promoting the p62/SQSTM1-LC3 interaction.

We have shown that our assay approach, which uses a non-saturating concentration of BafA, can robustly discriminate between autophagy stimulators and activators and offers a superior assay window compared to the use of saturating concentrations of BafA or CQ. Based on our results, we recommend that potential autophagy flux modulators should be tested in multiple concentrations and both in the presence as well as in the absence of BafA. This allows the identification of compounds which also increase LC3-II levels in the absence of this late-stage inhibitor, potentially indicating an inhibitory effect on autophagy. Third, to further confirm an inhibitory effect, the increase in p62/SQSMT1 in the absence of BafA can serve as a further indication for an autophagy flux inhibitor. In contrast, stimulators are expected to induce additional increase of the LC3-II levels on top of the change induced by the non-saturating concentrations of BafA, and to a significantly higher extent than in the absence of BafA. Autophagy stimulators in this setting should not increase p62/SQSTM levels in the absence of BafA.

## Materials and Methods

### Cell culture and compound treatment

HeLa cells were cultivated in RPMI 1640 (Gibco) supplemented with 1 mM sodium pyruvate (Gibco), 10 % FBS (Sigma Aldrich) and 50 μg/ml Gentamicin (Gibco) and passaged 2-3 times per week until they reached ~ 90 % confluency for up to 25 passage numbers. SK-N-MC cells were cultivated in MEM (Gibco) supplemented with 2 mM L-glutamine, 1 x MEM non-essential amino acids (Gibco), 10 % FBS and 50 μg/ml Gentamicin. For compound treatment and starvation experiments, cells were seeded on poly-D-lysine coated 6-well plates (culture area: 9.6 cm^2^) with a density of 30.000 cells / cm^2^ 24 h prior to cell treatment. On the day of treatment, a complete medium change was performed, switching to fresh medium containing compounds in the appropriate concentrations. The DMSO concentration was kept constant at 1 % in all compound treatment conditions and vehicle controls. DMSO stocks were prepared from rapamycin (ChemShuttle), bafilomycin A1 (Enzo), concanamycin A (Cayman chemical), chloroquine diphosphate (Biovision) and AbbVie-internal EP300 inhibitor [38] and stored in aliquots at −80 °C. For HBSS-starvation, HeLa cells were kept in HBSS (Gibco) for 3 h. For FBS-starvation, HeLa cells were cultivated in RPMI medium containing sodium pyruvate, gentamicin and 1.5 % FBS for 4 or 24 h. For harvesting, cells were washed once in PBS and scraped in phosphosafe extraction reagent (Merck Millipore) supplemented with 1x protease inhibitor cocktail (Roche).

### SDS-PAGE and Western blotting

Cell lysates in phosphosafe extraction reagent were sonicated for 15 sec (Amplitude: 35 %, Bandelin Sonoplus). Protein concentration was determined with a BCA Protein-Assay Kit (Thermo Fisher), applying BSA as a protein standard. For SDS-PAGE, 10 μg total protein per lane in Novex NuPAGE LDS-sample buffer supplemented with DTT were loaded on NuPAGE Novex 4-12 % bis-tris proteins gels in NuPAGE MES SDS running buffer (all: Thermo Scientific). Gels were blotted onto nitrocellulose membranes (Biorad) with a Biorad Trans-Blot Turbo Semidry blotting system for 30 min. After transfer, membranes were blocked in 5% milk (Sigma Aldrich) in TBS (Biorad) supplemented with 1% Tween-20 (Sigma Aldrich; TBS-T) for 1h. After washing membranes in TBS-T, primary antibodies (1:1.000 dilution in TBS-T) were incubated over night at 4°C. The following primary antibodies have been used: anti-LC3B antibody produced in rabbit (L7543, Sigma Aldrich), anti-SQSTM1/p62 antibody produced in mouse (ab56416, abcam), anti-actin (20-33) antibody produced in rabbit (A5060, Sigma Aldrich), GAPDH loading control monoclonal antibody (GA1R) produced in mouse (Thermo, MA5-15738), p-S6 ribosomal protein (S240/244) XP® rabbit mAB (5364S, Cell Signaling), S6 ribosomal protein mouse mAB (2317S, Cell Signaling), phospho-p70 S6 kinase (Thr389) antibody (9205, Cell Signaling), p70 S6 Kinase (49D7) rabbit mAb (2708, Cell Signaling). Membranes were washed with TBS-T and incubated with horseradish peroxidase-coupled secondary antibodies (1:5.000 dilution in TBS-T) for 90 min. Afterwards, membranes were washed and developed with a Chemidoc instrument (Biorad) applying SuperSignal West Pico PLUS Chemiluminescent Substrate or SuperSignal West Femto maximum sensitivity substrate (both: Thermo Fisher). Detection time of the instrument was set ensuring the longest possible exposure time before overexposure of the strongest band of interest occurred.

### MSD ELISA

To determine the ratio of Ser2488-phosphorylated to total mTOR, a MSD Multi-Spot Assay System (Phospho(Ser2448)/Total mTOR Assay Whole Cell Lysate Kit (K15170D9)) was applied according to the manufacturer’s protocol. Cell lysates were applied at 7.5 and 3.75 μg total protein concentration to ensure that the experiment ran within the linear range of the assay. For further data display, values obtained with 7.5 μg total protein were analyzed. Readouts were performed with a SECTOR Imager 6000 (Meso Scale Discovery).

### Data analysis and visualisation

Western blot band intensities were quantified using Fiji/Image J. Graphs and statistical analysis were prepared using GraphPad Prism 8.

## Acknowledgements

We thank Dr. Zhiqin Ji (AbbVie) for the synthesis of the AbbVie-internal EP300 inhibitor, Dr. Kenneth Bromberg (AbbVie) for help and support regarding activities on EP300-inhibitors, as well as Dr. Janina Ried (AbbVie) for advice on statistical analysis.

