## Supplementary information for "Creating a bottleneck: Robust LC3B lipidation analysis by adjusting autophagic flux with low concentrations of Bafilomycin A1"

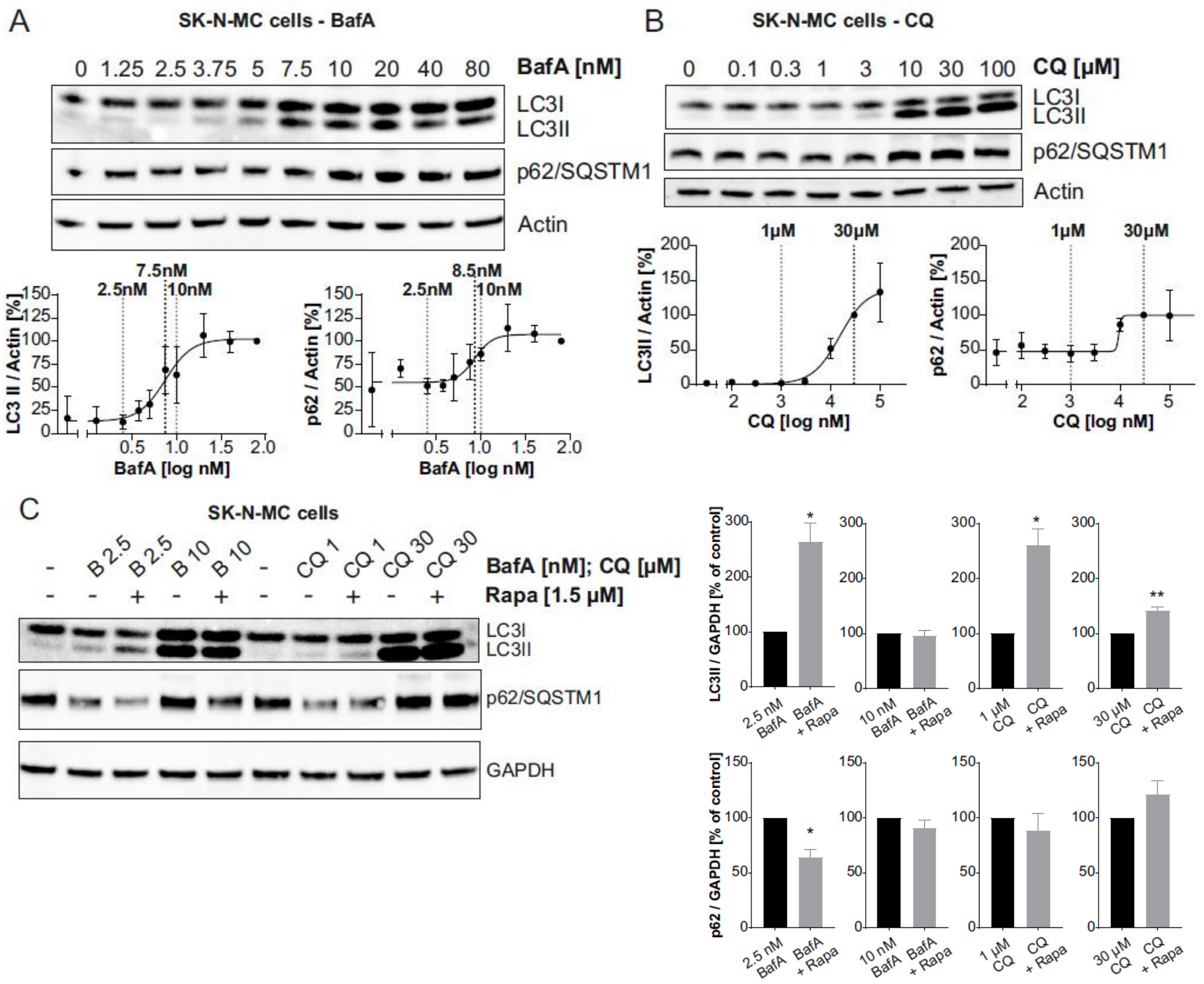


**Supplementary Figure 1. Applying non-saturating and saturating concentrations of late-stage autophagy inhibitors to SK-N-MC cells.** (**A**) SK-N-MC cells were treated with a concentration range of BafA (1.25 nM – 80 nM) for 24 h. Shown is a representative Western blot for LC3, p62/SQSMT1 and actin (top) as well as quantification based on Western blot analysis from 3 independent biological replicates (n = 3; mean +/- SEM). BafA inhibited the basal autophagic flux with an $EC_{50}$ of 7.5 and 8.5 nM based on the increases of LC3-II and p62/SQSTM1, respectively. For curve display, LC3-II or p62/SQSTM1 levels at 80 nM BafA were set to 100 %. For curve fitting, a non-linear regression curve fit (log(inhibitor) vs. response- variable slope (four parameters)) was applied. (**B**) SK-N-MC cells were treated with a concentration range (0.1 – 100 µM) of CQ for 24 h. Shown is a representative Western blot for LC3, p62/SQSMT1 and actin (top) as well as quantification based on Western blot analysis from 3 independent biological replicates (n = 3; mean +/- SEM). The calculated $EC_{50}$s with respect to LC3-II increase as well as p62/SQSTM1 accumulation were 15 µM and 9 µM, respectively. Curve calculation was done based on 3 independent biological replicates (mean +/- SEM). For curve display, LC3-II or p62/SQSTM1 levels at 30 µM CQ were set to 100 %. For curve fitting, a non-linear regression curve fit (log(inhibitor) vs. response- variable slope (four parameters)) was applied. **(C)** SK-N-MC cells were cotreated with either 2.5 nM or 10 nM BafA or either 1 µM or 30 µM CQ and 1.5 µM Rapamycin for 24 h. Shown is a representative Western blot for LC3, p62/SQSTM1 and GAPDH (left) as well as the quantification (right) from Western blots from 4 independent biological replicates (n = 4; mean +/- SEM). Statistics: One-sample t-test; compared to 100 % control (late-stage autophagy inhibitor only control, from left to right: 2.5 nM BafA, 10 nM BafA, 1 µM CQ, 30 µM CQ).

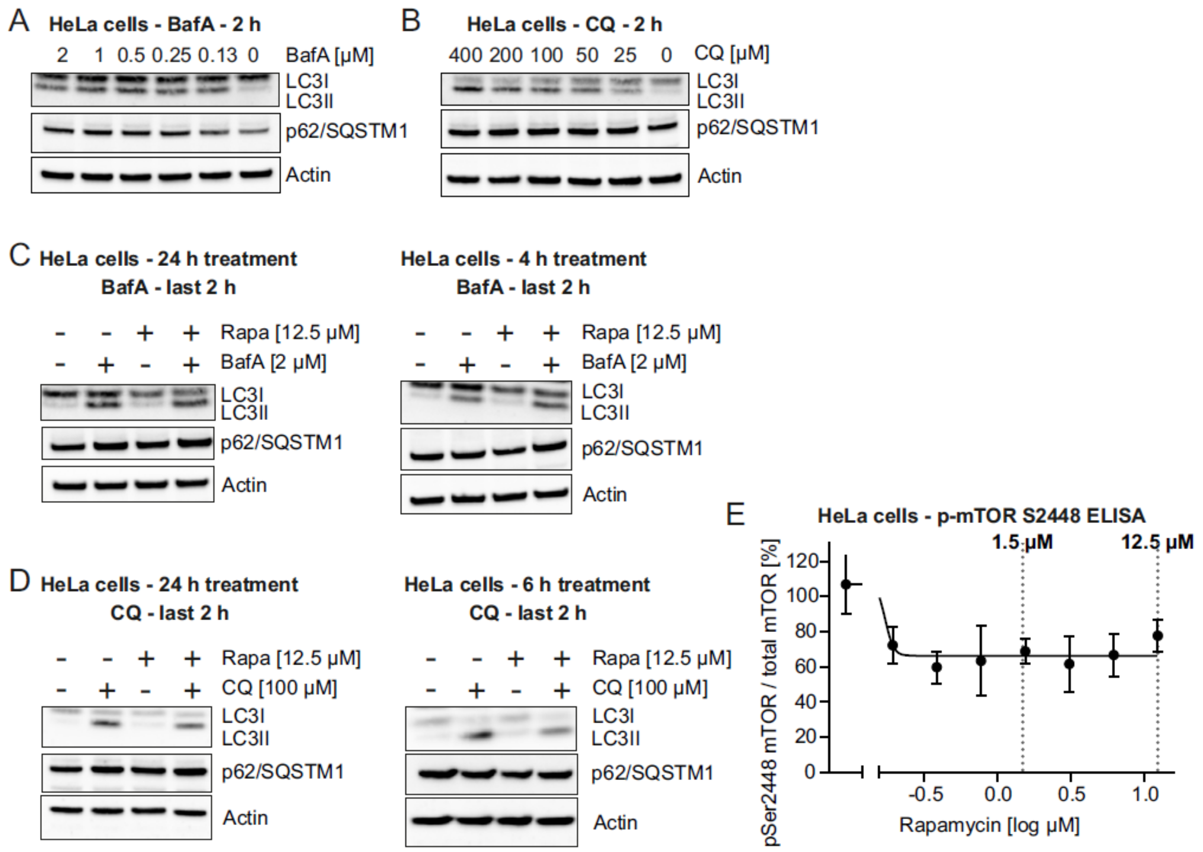


**Supplementary Figure 2. Short-time treatments of HeLa cells with late-stage autophagy inhibitors and mTOR ELISA assay.** For all experiments, representative Western blots for LC3, p62/SQSTM1 and actin are shown. **(A)** HeLa cells were treated with a concentration range (130 nM – 2 µM) of BafA for 2 h to determine a saturating concentration based on the LC3-II and p62/SQSTM1 increase. **(B)** HeLa cells were treated with a concentration range (25 µM – 400 µM) of CQ for 2 h to determine a saturating concentration based on the LC3-II and p62/SQSTM1 increase. **(C)** HeLa cells were treated with rapamycin for either 24 (left) or 4 h (right). BafA at a saturating concentration (2 µM) was added during the last 2 h of incubation with rapamycin. **(D)** HeLa cells were treated with rapamycin for either 24 (left) or 6 h (right). A saturating concentration of CQ (100 µM) was added during the last 2 h of incubation with rapamycin. **(E)** Lysates from HeLa cells treated with a dose range of rapamycin (200 nM – 12.5 µM) for 24 h were analyzed with a mTOR MSD ELISA assay multiplexing total mTOR and pS2448 mTOR. A maximum reduction of mTOR phosphorylation by 30-40 % was observed. The curve represents one biological replicate with 3 technical replicates (n=1; technical mean +/- SEM). For curve fitting, a nonlinear regression curve fit was applied (sigmoidal, four parameter logistic, X is log(concentration)).

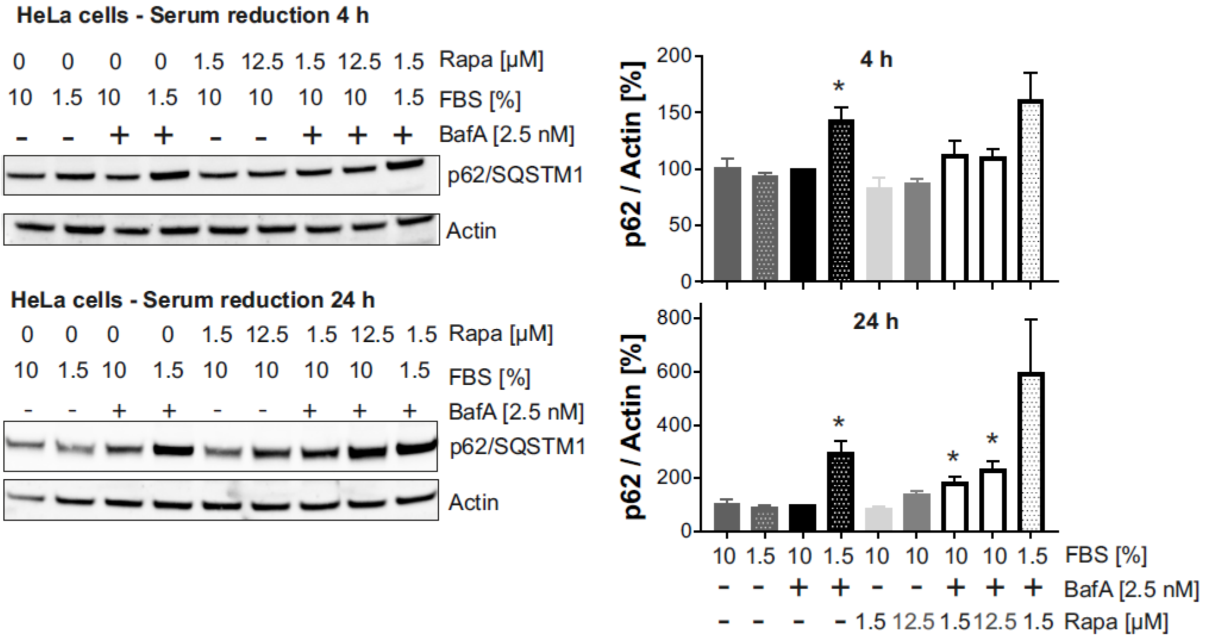


**Supplementary Figure 3. Capturing autophagic flux increase through serum starvation by applying a non-saturating concentration of BafA.** HeLa cells were kept in standard medium (10 % FBS) or in FBS-reduced medium (1.5 %) for 4 or 24 h. Medium change was combined with (co)-treatment of cells with 2.5 nM BafA and / or 1.5 or 12.5 µM rapamycin. Shown are representative Western blots of p62/SQSTM1 and actin (top) as well as quantifications (n=3, mean +/- SEM; bottom). Statistics: One-sample T-test; compared to 100 % control (DMSO, 10% FBS).
